# Pneumococcal phenotype and interaction with nontypeable *Haemophilus influenzae* as determinants of otitis media progression

**DOI:** 10.1101/200733

**Authors:** Joseph A. Lewnard, Noga Givon-Lavi, Paula A. Tähtinen, Ron Dagan

## Abstract

**Background:** All-cause otitis media (OM) incidence has declined in numerous settings following introduction of pneumococcal conjugate vaccines (PCVs) despite increases in carriage of non-vaccine pneumococcal serotypes escaping immune pressure. To understand the basis for declining incidence, we assessed the intrinsic capacity of pneumococcal serotypes to cause OM independently and in polymicrobial infections involving nontypeable *Haemophilus influenzae* (NTHi) using samples obtained from middle ear fluid and nasopharyngeal cultures before PCV7/13 rollout.

**Methods:** Data included OM episodes (11,811) submitted for cultures during a 10-year prospective study in southern Israel and nasopharyngeal samples (1588) from unvaccinated asymptomatic children in the same population. We compared pneumococcal serotype diversity across carriage and disease isolates with and without NTHi co-isolation. We also measured associations between pneumococcal phenotype and rate of progression from colonization to OM in the presence and absence of NTHi.

**Results:** Whereas pneumococcal serotype diversity in single-species OM is lower than in single-species colonization, serotype diversity does not differ significantly between colonization and OM in mixed-species episodes. Serotypes differed roughly 100-fold in progression rates, and these differences were attenuated in polymicrobial episodes. Vaccine-serotype pneumococci had higher rates of progression than non-vaccine serotypes. While serotype invasiveness was a weak predictor of OM progression rate, efficient capsular metabolic properties—traditionally thought to serve as an advantage in colonization—predicted an enhanced rate of progression to complex OM.

**Conclusions:** The lower capacity of non-vaccine serotypes to cause OM may partially account for reductions in all-cause OM incidence despite serotype replacement in carriage following rollout of PCVs.

## INTRODUCTION

Otitis media (OM) has historically been the leading cause of healthcare visits, antimicrobial prescribing, and surgical procedures among young children in high-income countries (1). *Streptococcus pneumoniae* (Spn) and nontypeable *Haemophilus influenzae* (NTHi) are the pathogens most commonly isolated from middle-ear fluid (MEF) in OM. Commensal carriage of these and other bacterial species in the upper-respiratory tract is the reservoir for middle-ear infection. Clinical severity ranges from acute or even asymptomatic presentations to complex OM (e.g., recurrent, nonresponsive, spontaneously-draining, or chronic OM and OM with effusion).

Because nearly all children carry Spn and NTHi in the first years of life, factors influencing progression from colonization to OM are ideal targets for treating or preventing disease. Pneumococcal conjugate vaccines (PCVs) with capsular antigens from 7, 10, and 13 Spn serotypes have recently been introduced to pediatric immunization schedules of most countries. Limited data from pre-licensure studies has contributed to uncertainty about the basis of PCV-mediated protection against OM (2–4), hampering efforts to interpret the considerable reductions reported in all-cause OM burden following PCV introduction in numerous settings (5–8) amid co-occurring changes in the circulation of vaccine-targeted and non-vaccine serotypes (9–11). Pneumococcal OM frequently involves the formation of polymicrobial biofilms with NTHi, and mixed-species infections represent a distinct clinical entity associated with recurrence, chronicity, and a unique serotype repertoire (12). However, the contribution of bacterial factors to disease progression remains poorly understood. Insight into this aspect of OM pathogenesis can guide interpretations of vaccine impact, and can inform serotype selection for extended-valency conjugate vaccines (13) as well as next-generation vaccine development (14, 15).

We analyzed isolates from MEF and asymptomatic nasopharyngeal colonization to better understand bacterial determinants of progression to complex OM. Using epidemiological surveillance data from southern Israel prior to PCV7/13 introduction, we compared pneumococcal serotype distributions of single-species and mixed-species carriage and OM, and measured pathogen-specific rates of OM progression from asymptomatic colonization. We identified roughly 100-fold differences in rates of pneumococcal serotype-specific progression to complex OM, and found that these differences were attenuated in mixed-species episodes involving NTHi—further signaled by enhanced serotype diversity in polymicrobial OM relative to single-species episodes. Pneumococcal serotypes targeted by PCV13 are among the most virulent in terms of OM progression, a finding that may in part account for reductions in all-cause OM incidence following PCV introduction despite serotype replacement in pneumococcal carriage.

## RESULTS

### Study enrollment

Data came from several studies undertaken prior to PCV7 introduction in southern Israel; we included samples obtained before July, 2008 in analyses (**Table 1**). Nasopharyngeal carriage of Spn and NTHi was monitored among PCV7/13-unvaccinated children ages 2 to 18 months old enrolled in a randomized trial of PCV7 dosing strategies between 2005 and 2008 (16, 17). In total, children submitted 1588 samples, from which Spn was detected in 743 (46.8%) swabs; NTHi was detected in 524 (33.0%) swabs, and the two species were co-isolated in 376 (23.7%). A ten-year prospective study of the incidence of severe OM cases necessitating MEF culture (as indicated by complex manifestations; detailed in **Materials and Methods**) supplied 11,811 cases (12), with 4165 (35.3%) positive for Spn, 4813 (40.8%) positive for NTHi, and 1589 (13.5%) positive for the two species. Other previously-conducted laboratory and epidemiological studies supplied phenotype information about pneumococcal serotypes (18–21). Further descriptive details including age-specific OM incidence and carriage prevalence, and serotype frequencies in carriage and disease, are included as supporting information (**Figure S1**, **Table S1**).

**Table 1:** Data sources

| Variables measured | Measurement details | Coverage | Citation |
| --- | --- | --- | --- |
| Pneumococcal OM incidence | All episodes of OM necessitating MEF culture from children <18 months old presenting for care at Soroka University Medical Center between July 1999 and June 2008 | 11811 episodes, with 9397 Spn or NTHi positive | (26) |
| Nasopharyngeal pneumococcal carriage prevalence | Unvaccinated Bedouin and Jewish children sampled at scheduled visits, ages 2-18 months, enrolled in a PCV7 dosing study | 769 children submitting 1588 swabs, with 891 Spn or NTHi positive | (17) |
| Resistance to neutrophil-mediated killing<sup>1</sup> | Proportion surviving a complement-independent *in vitro* surface killing assay for each pneumococcal serotype | 14 serotypes<sup>2</sup> | (18) |
| Magnitude of anionic surface charge<sup>1</sup> | (Negative) zeta potential of fixed-density suspension of pneumococcal serotype in phosphate-buffered saline | 48 serotypes<sup>2</sup> | (19) |
| Capsular size<sup>1</sup> | Zone of exclusion of fluorescent dextran molecules around pneumococcal serotype diplococcus | 15 serotypes<sup>2</sup> | (18) |
| IPD case-fatality ratio | Serotype-specific 30-d mortality during IPD | 37 serotypes<sup>2</sup> | (20) |
| Metabolic efficiency | Inverse of number of carbons per capsular polysaccharide repeat unit of pneumococcal serotype | 54 serotypes<sup>2</sup> | (18) |
| Invasiveness | Proportion of carriage events leading to IPD | 36 serotypes<sup>2</sup> | (21) |

<sup>1</sup>*In vitro* measurements of serotype properties were obtained from isogenic capsular-switch mutants
<sup>2</sup>We list serotypes for which phenotypic data were collected in **Table S1**.

### Pneumococcal serotype distribution in single-species and mixed-species colonization and otitis media

If serotype factors play a role in progression of pneumococcal colonization to OM, the diversity of serotypes isolated from MEF would be expected to differ from the diversity of serotypes carried in the nasopharynx (22). To investigate this hypothesis, we calculated the Simpson’s Diversity Index (*D*) for pneumococcal serotypes isolated from MEF and from carriage, with and without co-occurring NTHi, and tested for differences in serotype diversity (**Figure 1**); *D* measures the probability for any two randomly-chosen isolates to belong to different serotypes (see **Materials and Methods**).

**Figure 1:**
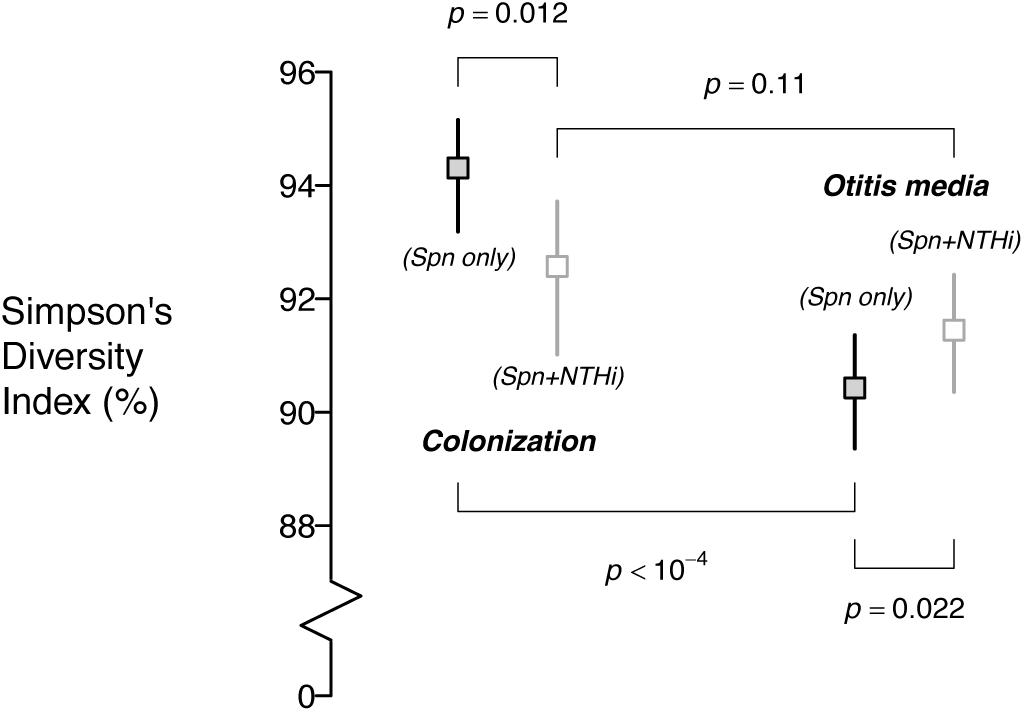
Pneumococcal serotype diversity in carriage and MEF isolates, with and without co-occurring NTHi. Pneumococcal serotype diversity is lower in single-species OM than single-species colonization, suggesting a role of serotype factors in progression. This difference is not apparent in mixed-species colonization and OM. Lines denote 95% confidence intervals around estimates.

Pneumococcal serotype diversity was higher in nasopharyngeal isolates than in MEF isolates both in the presence and in the absence of NTHi (*p*<10^−4^), consistent with the hypothesis that a limited number of serotypes cause disproportionate disease burden. However, in mixed infections, the difference was no longer significant (*p*=0.11). Whereas serotype diversity was lower in the presence of NTHi during colonization, an opposite relationship was seen in MEF isolates, where diversity was significantly lower in single-species than in mixed-species infections (*p*=0.022).

We next sought to determine whether similar pneumococcal serotype diversity in mixed-species colonization and OM owed to the isolation of serotypes from nasopharyngeal and MEF samples at similar frequencies. We used Kullback-Leibler divergence to measure the relatedness of serotype distributions from single-species OM, single-species carriage, and mixed-species carriage to the serotype distribution of mixed-species OM (**Figure 2**). We identified lower divergence between serotype distributions of mixed-species carriage and OM than between serotype distributions of single-species carriage and mixed-species OM (*p*<0.01)—a finding that supports serotype-specific polymicrobial interactions in carriage as the basis for patterns of cooccurrence in disease. Divergence between the pneumococcal serotype distributions of mixed-species OM and single-species OM was lower, however, suggesting a overriding role of pneumococcal serotype in both single-species and mixed-species OM progression.

**Figure 2:**
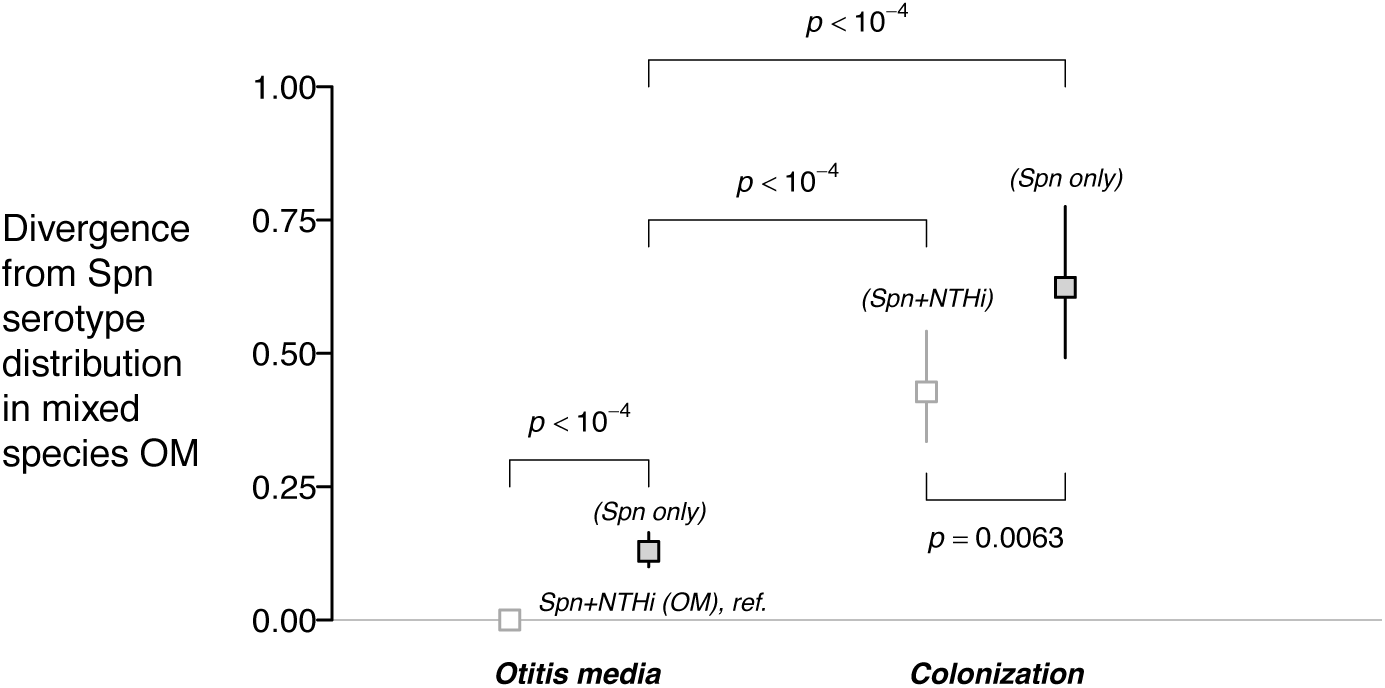
Divergence of pneumococcal serotype distribution in single-species and mixed-species carriage and OM. To determine whether serotype-specific interactions of *S. pneumoniae* with NTHi contribute to the pneumococcal serotype distribution of mixed-species OM episodes, we calculated Kullback-Leibler divergence in pneumococcal serotype distributions from that of mixed-species OM episodes for: single-species (Spn) OM, mixed-species carriage (Spn+NTHi), and single-species carriage. A value of zero indicates an exact match with the pneumococcal serotype distribution of mixed-species OM episodes; increasing values reflect greater divergence. Lines denote 95% credible intervals around estimates.

### Determinants of progression from colonization to OM

We next compared pathogen-specific rates of progression to OM episodes necessitating MEF in order to identify bacterial factors associated with virulence. We derived the odds ratio of disease as an expression for the relative rate of progression from asymptomatic colonization to OM (see **Materials and Methods**).

Progression rates for pneumococcal serotypes spanned over two orders of magnitude (**Figure 3**). The highest progression rates were detected among PCV13 serotypes, including 1, 3, 5, and 7F; for the latter two serotypes, which were detected in 24 and 101 single-species OM cases respectively, no instances of single-species carriage were detected. Although we also detected no single-species carriage of 7B and 24F, these serotypes accounted for only 6 and 4 single-species OM episodes, respectively. Though not detected in mixed-species colonization, serotypes 8, 33F, and 15B/C were isolated from 4, 28, and 48 instances of mixed-species OM, respectively. Unencapsulated pneumococci had the lowest measured progression rates for single-species OM, while no instances of single-species OM involved serotypes 22F or 19B. We did not identify strong statistical evidence (defined by a 95% confidence interval entirely above one) that any serotype had a higher progression rate than NTHi.

**Figure 3:**
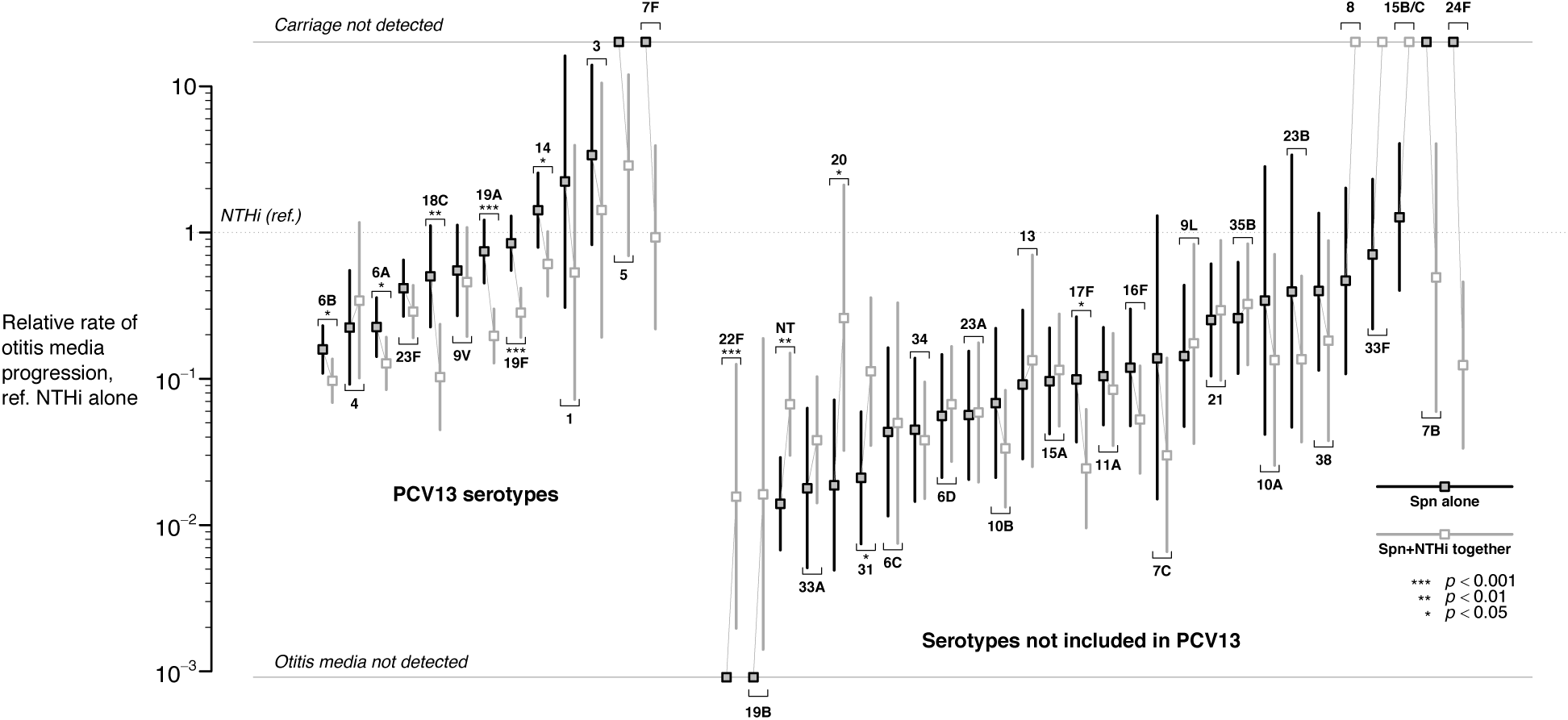
Pneumococcal serotype associations with otitis media progression. The rate ratio of OM progression from colonization (as measured by the adjusted odds ratio; see Methods for derivation) is calculated for each *S. pneumoniae* serotype, in single-species and mixed-species episodes, relative to NTHi. Analyses control for season (Dec-Mar or other) and Jewish/Bedouin ethnicity. Unencapsulated serotypes are labeled “NT”. Models are estimated with an exchangeable correlation structure to account for repeated sampling of individual children in the carriage data. Lines denote 95% confidence intervals around estimates.

Serotype differences in progression for mixed-species episodes were attenuated in comparison to differences in progression for single-species episodes. We identified lower progression rates in mixed-species episodes involving serotypes that tended otherwise to be virulent when colonizing in the absence of NTHi, including 6A, 6B, 14, 18C, 17F, 19A, 19F, and 31. Only nontypeable pneumococci and serotypes 20 and 31 showed increased progression rates in association with NTHi.

To understand this variation in progression to OM, we used regression models to calculate associations between phenotypes and progression rates for single-species and mixed-species episodes. To enable comparison of effect sizes, we scaled all phenotype variables to unit variance (**Table 2**; **Table S2**). We estimated a 50% (95%CI: 31-72%) increase in progression rate for each increase by one standard deviation in the metabolic efficiency of capsule production—a strong predictor of pneumococcal fitness in colonization (18) and capacity to co-colonize with NTHi (22)—which was not apparent in polymicrobial episodes. We identified a weak, 14% (1-29%) increase in progression rate for each increase by one standard deviation in serotype invasiveness, reflecting our findings that invasive disease-associated serotypes (e.g., 1, 4, 5, 14) were among those more likely to cause OM together with serotypes such as 3 and 15B/C, which are less invasive. We identified similar associations between higher progression rate and thinner encapsulation and lower case-fatality ratios, which are associated with invasiveness (23), and with a weaker surface charge in mixed-species episodes.

**Table 2:** Pneumococcal phenotype associations with otitis media progression

| Pneumococcal attribute<sup>1</sup> | Relative progression rate, per 1SD increase in covariate (95% CI)<sup>2</sup> | Adjusted relative progression, per 1SD increase in covariate (95% CI)<sup>2,3</sup> | Adjusted relative increase in progression rate, per 1SD increase in covariate, for Spn+NTHi (95% CI) |
| --- | --- | --- | --- |
| Metabolic efficiency (18) |  |  |  |
| Spn only | **1.51 (1.32, 1.74)** | **1.50 (1.31, 1.72)** | ref. |
| Spn+NTHi | 1.09 (0.96, 1.25) | 1.07 (0.94, 1.22) | **0.72 (0.59, 0.86)** |
| Surface anionic charge (19) |  |  |  |
| Spn only | 1.02 (0.95, 1.10) | 1.03 (0.95, 1.11) | ref. |
| Spn+NTHi | **0.80 (0.72, 0.89)** | **0.79 (0.71, 0.88)** | **0.77 (0.68, 0.88)** |
| Resistance to neutrophil-mediated killing (18) |  |  |  |
| Spn only | 1.05 (0.93, 1.19) | 1.04 (0.92, 1.16) | ref. |
| Spn+NTHi | 1.07 (0.94, 1.21) | 1.05 (0.93, 1.20) | 1.02 (0.85, 1.21) |
| Capsule width (18) |  |  |  |
| Spn only | **0.87 (0.78, 0.98)** | **0.87 (0.77, 0.97)** | ref. |
| Spn+NTHi | **0.81 (0.72, 0.91)** | **0.80 (0.71, 0.90)** | 0.93 (0.78, 1.09) |
| Case-fatality ratio in IPD (20) |  |  |  |
| Spn only | **0.71 (0.63, 0.81)** | **0.71 (0.63, 0.81)** | ref. |
| Spn+NTHi | **0.73 (0.64, 0.84)** | **0.74 (0.65, 0.85)** | 1.04 (0.86, 1.25) |
| Invasiveness (IPD per carriage episode) (21) |  |  |  |
| Spn only | **1.15 (1.02, 1.30)** | **1.14 (1.01, 1.29)** | ref. |
| Spn+NTHi | 1.12 (0.99, 1.26) | 1.11 (0.99, 1.25) | 0.97 (0.82, 1.15) |

<sup>1</sup>Pneumococcal attributes are defined in Table 1.
<sup>2</sup>Progression rates are compared via the odds ratio, as calculated by logistic regression with an exchangeable correlation structure to account for repeated carriage observations from individual children. Due to collinearity among serotype measurements, individual models are fitted for each association. Our derivation of the odds ratio as the relative progression rate is in the Methods.
<sup>3</sup>Models control for Jewish/Bedouin ethnicity and respiratory virus season (December to March).

Our analyses also revealed a 77.5% (46.3%, 114.5%) higher progression rate during the months with the highest degree of respiratory virus transmission (December to March), compounding elevated carriage prevalence during these months as a driver of seasonal OM incidence (17).

## DISCUSSION

We sought to better understand the progression of Spn and NTHi to complex OM manifestations using prospectively-gathered epidemiological surveillance data from Israel. We identified over 100-fold differences in the rate at which pneumococcal serotypes progress from colonization to OM. Serotypes 1, 3, 5, 7B, 7F, and 24F showed the greatest virulence as causes of single-species infections; in addition, serotypes 8, 15B/C, and 33F showed particularly high rates of mixed-species progression with NTHi. A narrower range of mixed-species progression rates, together with elevated serotype diversity in mixed-species disease relative to single-species disease, suggested interactions with NTHi may attenuate serotype-specific differences in disease potential. While we identified a weak positive association between OM progression rates and risk for serotypes to cause invasive pneumococcal disease, we also found that each standard-deviation increase in the metabolic efficiency of a serotype—typically a marker for colonization fitness and lower virulence (18)—was associated with a 50% increase in OM progression rate. Taken together, our findings substantiate the role of bacterial factors, including serotype-specific interactions with NTHi, in the pathogenesis of complex pneumococcal OM.

Our findings aid interpretation of PCV impact against OM. Shortly after vaccine introduction, concerns arose suggesting the replacement of PCV-targeted serotypes by non-vaccine serotypes in carriage might offset vaccine impact against OM (24), consistent with observations in IPD (25). Nonetheless, studies in Israel (26) and other settings (5) have reported declining all-cause OM incidence following PCV rollout. Although comparable epidemiological surveillance is not undertaken in the United States, reductions in all-cause OM incidence relative to the pre-vaccine era (27) are apparent from the proportion of children experiencing primary and subsequent OM episodes during each year of life (28, 29), and from reduced healthcare utilization for severe OM (8, 30–32). While the effects of serotype replacement in carriage are apparent from the changing serotype distribution of OM episodes in Israel as well as in US studies (26, 28, 33), our finding of lower virulence among non-vaccine replacement serotypes helps to account for overall reductions in OM incidence following vaccine introduction. Because tissue damage sustained during early-life OM episodes historically associated with PCV13 serotypes contributes to risk for secondary infections (34), PCV7/13-mediated protection against OM caused by these virulent lineages may also contribute indirectly to reduced incidence of OM caused by other pathogens, including NTHi and non-vaccine pneumococcal serotypes (26, 35).

Previous studies demonstrating differences in pneumococcal serotype-specific disease potential have provided important evidence of the role of the capsule as a virulence factor in OM (24, 36) and other pneumococcal disease manifestations (21, 37). Smaller sample sizes, however, limited the statistical power of such studies and thus their ability to detect differences in serotype-specific rates of progression, contributing to concerns that serotype replacement may offset vaccine-attributable reductions in OM burden (24). Data from over 13,000 isolates gathered through prospective, population-based surveillance enabled us to compare progression rates of 41 serotypes, stratifying according to co-isolation with NTHi and thus identifying greater differences in disease potential than were previously recognized.

The biological basis for variation among serotypes in the capacity to cause OM is incompletely understood and likely multifactorial. Epidemiological observations suggest OM is frequently a consequence of prior viral infection in the upper respiratory tract (29, 39, 40), a finding supported by *in vivo* studies demonstrating that altered expression of bacterial adherence receptors and inflammation after viral infection facilitate bacterial infiltration of the middle ear (38). A growing body of evidence suggests such bacterial-viral interactions behind disease progression may be serotype-specific. In experimentally-challenged ferrets, influenza A infection enhanced the recovery of Spn in nasal wash in a serotype-dependent manner, and disproportionately enhanced the capacity of serotype 7F to cause OM (41); this serotype likewise showed considerable disease potential in our study. In mice, influenza A infection modified proinflammatory cytokine responses to Spn differentially among serotypes, and was prerequisite to pathogenesis following nonlethal challenge with serotypes 19F and 7F (42). Although the immunological pathway was not determined, a separate study showed that live attenuated influenza vaccination triggered increases in the duration and density of upper respiratory carriage of these serotypes, which may contribute to enhanced risk of upper respiratory mucosal disease (43). Respiratory syncytial virus (RSV) is the most prevalent virus in MEF (44) and may contribute to OM pathogenesis through similar dysregulation of local anti-pneumococcal immune responses (38). Additionally, *in vitro* studies have demonstrated serotype-dependent enhancement of pneumococcal adherence to RSV-infected epithelial cells (45).

While interactions with NTHi including biofilm formation may alter species-specific viral-bacterial interactions (46), our analysis suggests that the rank order of serotype virulence is nonetheless relatively consistent in the presence and absence of NTHi. Given evidence from our study of capsule as a virulence determinant, the attenuation we observe in serotype-specific virulence differences in association with NTHi, and enhanced serotype diversity in mixed-species OM, may relate to down-regulation of capsule expression in mixed-species biofilms (47). Our finding that many serotypes show lower rates of progression to OM when co-colonizing with NTHi reflects “stable” characteristics of mixed-species infection in a chinchilla model (48). However, this observation in our study may also reflect acquisition of NTHi at older, less-susceptible ages (**Figure S1**), or reduced sensitivity for the detection of bacteria in polymicrobial biofilms (34, 49).

Our analysis has certain limitations. An ideal prospective study of OM progression would monitor nasopharyngeal carriage and OM incidence with MEF culture within a cohort of children to assess pathogen-specific OM incidence rates during colonization. Because the sample size needed to characterize serotype differences in progression would be prohibitively large in such a study, however, microbiologically-detailed prospective studies of carriage and disease provide a good alternative for characterizing pathogen-specific virulence (24, 35, 36, 50). The validity of evaluating progression via OM incidence per carrier is suggested by meta-analytic findings of high concordance (>80%) in Spn and NTHi detection from nasopharyngeal and MEF cultures in severe OM cases (51), such as those necessitating MEF culture in our study. Nonetheless, molecular diagnostic tools may offer enhanced sensitivity in comparison to culture as performed here (52–54). Such methods may be particularly important for estimating progression rates of serotypes that are rarely detected in carriage or disease: despite the large sample in our study, odds ratio estimates were sensitive to the lack of detection of episodes of either carriage (e.g., serotypes 5, 7B, 7F, 8, 15B/C, 24F, 33F) or disease (serotypes 19B, 22F) in either single-or mixed-species contexts. Although our analysis does not characterize the contribution of viral infections to disease onset and progression (39, 55), their impact is suggested in our analysis by elevated progression rates during the winter months.

It is important to note that our sample was enriched with complex OM cases due to the indications for obtaining MEF culture in the original prospective study. Thus, comparisons in our analysis reflect differences in progression to severe OM, and may not represent the broader spectrum of illness including acute OM episodes. Children who experience complex OM may differ from those experiencing acute OM; higher prevalence of pneumococcal, NTHi, and polymicrobial colonization has been reported among otitis-prone children as compared to non– otitis-prone children owing to deficient mucosal antibody responses (56, 57). High rates of NTHi progression in the absence of Spn may also reflect predominance of this pathogen in secondary infections following tissue damage associated with early-life OM (34). Nonetheless, the use of NTHi as a reference category does not impact our ability to compare relative progression rates among serotypes.

Using data obtained prior to the establishment of widespread immunity against certain pneumococcal serotypes through routine PCV7/13 immunization, our study demonstrates greater variation in the capacity of pneumococcal serotypes to cause OM than has been previously recognized, as well as serotype-dependent alteration of disease potential associated with interaction with NTHi. High OM progression rates for PCV-targeted serotypes help to account for observations of reduced all-cause OM incidence following vaccine rollout despite serotype replacement in carriage. Epidemiological evidence from our study should inform mechanistic studies addressing the biological basis for variation in the capacity of serotypes to progress to OM in single-species and polymicrobial episodes. In addition, our estimates of serotype-specific virulence can inform development of future anticapsular and next-generation vaccines.

## METHODS

### Setting

Previously-published studies provided data on incidence of severe OM necessitating MEF culture (26) and prevalence of bacterial carriage (16, 17) among Bedouin and Jewish children in the Negev region of southern Israel. The Bedouin population is transitioning from nomadic lifestyles to permanent settlements, and has larger family sizes, higher levels of overcrowding, and lower socioeconomic status in comparison to the nearby Jewish population (58). Bedouin children tend to experience higher rates of bacterial carriage and disease than Jewish children, despite receiving care from the same facilities (17).

### Datasets

#### OM incidence and carriage prevalence

Incidence of OM episodes necessitating MEF culture is monitored routinely at the Soroka University Medical Center (SUMC) through an ongoing prospective, population-based epidemiological surveillance program. Over 95% of children in the Negev region receive care at SUMC. Indications for MEF culture are based on clinical severity and include, but are not limited to, previous OM or tube insertion at any time, high-grade fever or toxic appearance, and spontaneous drainage, as detailed previously (26); these criteria have not changed during the study period.

Prevalence of nasopharyngeal Spn and NTHi carriage was monitored in a pre-implementation trial of PCV7 dosing schedules, wherein receipt of a first vaccine dose was randomized to between 2 and 18 months of age (16); data regarding pneumococcal and NTHi co-colonization have been published previously (17, 22). We included data from all visits by unvaccinated children up to age 18 months.

#### Bacteriological procedures

Samples of MEF were obtained from tympanocentesis or spontaneous drainage and placed in MW173 Amies transport medium (Transwab; Medical Wire and Equipment, Potley, UK) before being plated, within 16 hours, on trypticase agar (5% sheep blood and 5 μg/mL gentamicin) and chocolate agar media, followed by 48 hours’ incubation at 35°C in 5% CO_2_. Laboratory procedures for identification of Spn and NTHi were consistent in the studies of carriage and MEF, and have been described previously (16). Pneumococcal serotypes were determined by the Quellung reaction (antisera from Statens Serum Institut, Copenhagen, Denmark). Studies received ethics approval from SUMC, and secondary analyses were exempted by the institutional review board at Harvard TH Chan School of Public Health.

#### Pneumococcal serotype measurements

We used previously-obtained measurements of pneumococcal phenotypes in our analyses of factors associated with progression rate. These included the negative surface charge of the capsule (a determinant of susceptibility to phagocytosis) (19); the metabolic efficiency of capsule production, measured by the inverse of the number of carbons per repeat unit of the polysaccharide (18); the ability of serotypes to survive neutrophil-mediated killing in an *in vitro* surface assay (18); the width of the capsule (18); the likelihood for serotypes to cause death during invasive pneumococcal disease (IPD) (20); and the likelihood for serotypes to progress from carriage to IPD (21).

### Statistical analysis

#### Serotype distribution in carriage and otitis media

We measured serotype diversity (*D*) as

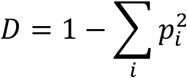

where *p*_*i*_ values indicated the proportion of isolates belonging to serotype *i*. We used bootstrap resampling to measure confidence intervals around estimates and to test for differences in diversity in carriage and OM, applying a cluster bootstrap of children to account for repeated sampling in the carriage studies (22).

We used Kullback-Leibler divergence to measure similarity of the pneumococcal serotype distribution in mixed-species OM to the serotype distributions of the following clinical entities: pneumococcal OM without NTHi, pneumococcal carriage without NTHi, and mixed-species carriage of Spn and NTHi. We sampled from Dirichlet-multinomial posterior distributions of serotype frequencies, applying a flat (Jeffreys) prior to account for uncertainty in sparse observations (59). Measures closer to zero indicated greater similarity to the pneumococcal serotype distribution of mixed-species OM. Consistent with our analyses of Simpson diversity, we generated credible intervals and conducted hypothesis testing via the bootstrap for disease isolates, and via the cluster bootstrap for carriage isolates.

#### Variation in otitis media progression rate

Odds ratios calculated from counts (*Y*) of pathogen-specific carriage and disease episodes supplied the relative rate of progression from colonization to OM, measured as (cases/year)/carrier. The theoretical basis of this interpretation is as follows: defining the prevalence of colonization by agent(s) *i* and *j* as *π*_*i*_ and *π*_*j*_, and the rates of OM incidence as *λ*_*i*_and *λ*_*j*_,

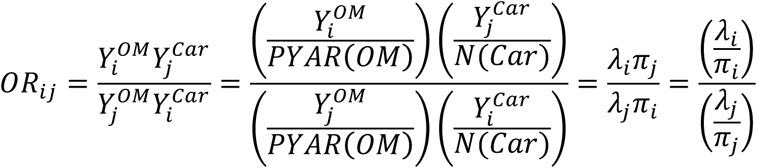
where *PYAR(OM)* and *N(Car)* refer to total person-years at risk for OM and children at risk for carriage, respectively, in the two datasets.

Based on the same premise, we used the odds ratio of OM to quantify the association between pneumococcal serotype factors and rate of progression from colonization to OM in the presence and absence of NTHi. Defining *e*^*α*_0_^ and *e*^*β*_0_^ as the “baseline” rate of OM incidence and prevalence of colonization, respectively, and *e*^*α*_1_X^ and *e*^*β*_1_X^ as the multiplicative impact of a serotype factor *X* on incidence and prevalence, respectively,

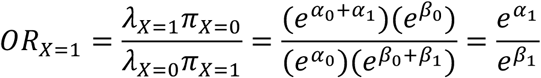
which again represents the relative progression rate as the fold increase in incidence for *X*=1 (ref. *X*=0), normalized by any effect of *X* on carriage prevalence.

We used logistic regression to calculate odds ratios and adjusted odds ratios, controlling for the seasonal peak in virus transmission (months December to March) and Jewish or Bedouin ethnicity. Because differing numbers of children were retained in the unvaccinated group of the carriage study across ages, it was not possible to adjust for age in the odds ratio formulation above; however, the narrow age range considered (≤18 months) limits bias. We fitted models via generalized estimating equations with an exchangeable correlation structure to account for repeated sampling of children in the carriage studies.

## ACKNOWLEDGMENTS

This work was supported by Pfizer (CP147216 to JAL). The original carriage studies were supported by grants from Wyeth/Pfizer and Berna/Crucell to RD. The funder had no role in study design, data collection and analysis, decision to publish, or preparation of the manuscript. The authors thank Marc Lipsitch for helpful comments.

JAL received research funds from Pfizer to Harvard University for the study (grant CP147216). JAL has also received consulting fees from Pfizer. RD has received grants and research support from Berna/Crucell, Wyeth/Pfizer, Merck, and Protea; has been a scientific consult for Berna/Crucell, GlaxoSmithKline, Novartis, Wyeth/Pfizer, Merck, and Protea; has been a speaker for Berna/Crucell, GlaxoSmithKline, and Wyeth/Pfizer; and is a shareholder of Protea/NASVAX. NG-L and PAT report no conflicts.

---

**Table S1:** Pneumococcal serotype frequencies, by source and NTHi co-isolation.

| Serotype | Middle ear fluid isolates |  | Nasopharyngeal isolates |  |
| --- | --- | --- | --- | --- |
|  | NTHi absent | NTHi present | NTHi absent | NTHi present |
| 1 | 45 | 13 | 1 | 1 |
| 3 | 136 | 29 | 2 | 1 |
| 4 | 28 | 25 | 6 | 3 |
| 5 | 100 | 21 | 0 | 2 |
| 6A | 125 | 100 | 27 | 36 |
| 6C | 5 | 2 | 5 | 2 |
| 6B | 163 | 130 | 47 | 62 |
| 7B | 6 | 5 | 0 | 1 |
| 7C | 3 | 3 | 1 | 4 |
| 7F | 24 | 17 | 0 | 2 |
| 8 | 16 | 4 | 2 | 0 |
| 9A | 3 | 6 | 2 | 1 |
| 9L | 13 | 7 | 4 | 2 |
| 9N | 2 | 2 | 5 | 2 |
| 9V | 104 | 58 | 9 | 6 |
| 10A | 7 | 5 | 1 | 2 |
| 10F | 2 | 4 | 3 | 0 |
| 11A | 22 | 16 | 10 | 9 |
| 12A | 8 | 1 | 0 | 0 |
| 12F | 18 | 11 | 0 | 0 |
| 13 | 7 | 6 | 4 | 2 |
| 14 | 379 | 231 | 13 | 18 |
| 15A | 18 | 17 | 9 | 7 |
| 15B/C | 38 | 48 | 3 | 0 |
| 16F | 19 | 12 | 8 | 11 |
| 17F | 13 | 7 | 6 | 13 |
| 18A | 4 | 5 | 0 | 0 |
| 18B | 1 | 1 | 1 | 0 |
| 18C | 74 | 20 | 7 | 9 |
| 19A | 304 | 135 | 20 | 31 |
| 19B | 0 | 1 | 2 | 1 |
| 19F | 458 | 228 | 26 | 37 |
| 20 | 3 | 6 | 6 | 1 |
| 21 | 31 | 26 | 6 | 4 |
| 22A | 0 | 2 | 0 | 0 |
| 22F | 0 | 1 | 2 | 1 |
| 23A | 8 | 7 | 7 | 6 |
| 23B | 8 | 8 | 1 | 3 |
| 23F | 230 | 206 | 26 | 33 |
| 24B | 2 | 2 | 0 | 0 |
| 24B | 2 | 2 | 0 | 0 |
| 24F | 4 | 5 | 0 | 3 |
| 31 | 5 | 10 | 11 | 4 |
| 33A | 3 | 7 | 7 | 7 |
| 33F | 19 | 28 | 3 | 0 |
| 34 | 6 | 8 | 6 | 10 |
| 35A | 1 | 2 | 0 | 0 |
| 35B | 34 | 37 | 6 | 5 |
| 35F | 4 | 0 | 6 | 0 |
| 37 | 3 | 2 | 1 | 0 |
| 38 | 22 | 9 | 3 | 2 |
| NT | 10 | 15 | 35 | 11 |

**Table S2:**
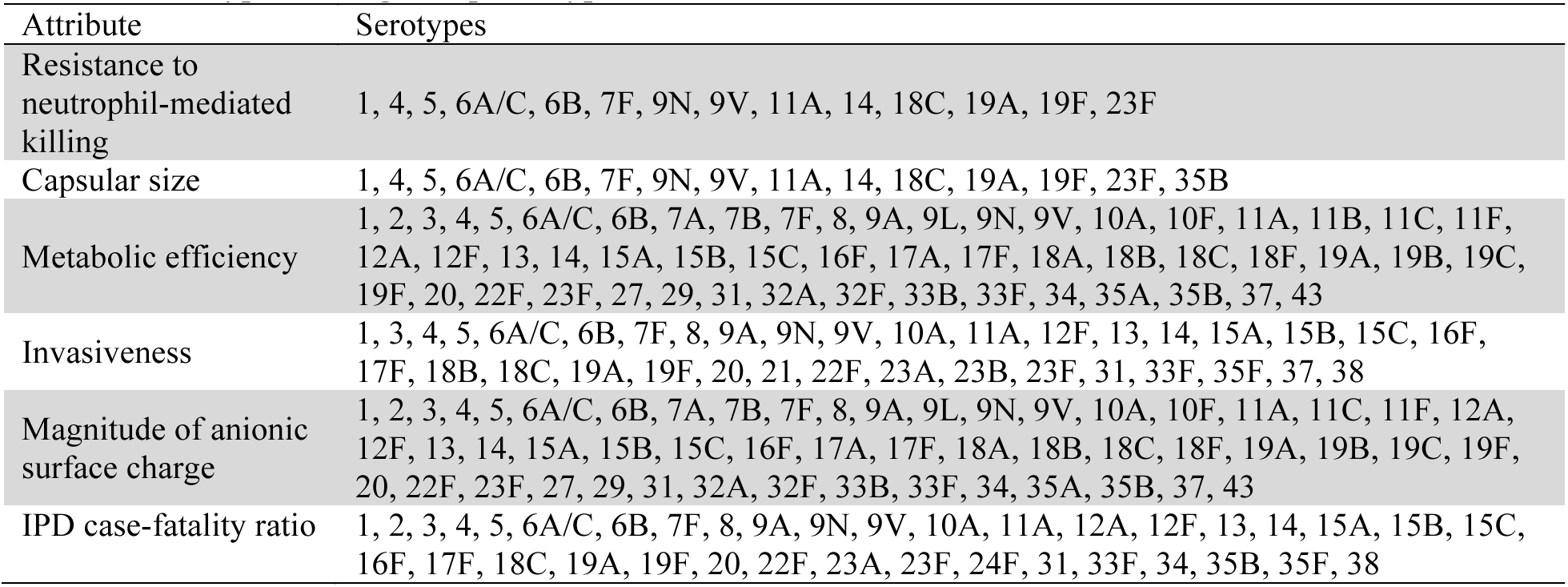
Serotype coverage for phenotypic measurements.

| Attribute | Serotypes |
| --- | --- |
| Resistance to neutrophil-mediated killing | 1, 4, 5, 6A/C, 6B, 7F, 9N, 9V, 11A, 14, 18C, 19A, 19F, 23F |
| Capsular size | 1, 4, 5, 6A/C, 6B, 7F, 9N, 9V, 11A, 14, 18C, 19A, 19F, 23F, 35B |
| Metabolic efficiency | 1, 2, 3, 4, 5, 6A/C, 6B, 7A, 7B, 7F, 8, 9A, 9L, 9N, 9V, 10A, 10F, 11A, 11B, 11C, 11F, 12A, 12F, 13, 14, 15A, 15B, 15C, 16F, 17A, 17F, 18A, 18B, 18C, 18F, 19A, 19B, 19C, 19F, 20, 22F, 23F, 27, 29, 31, 32A, 32F, 33B, 33F, 34, 35A, 35B, 37, 43 |
| Invasiveness | 1, 3, 4, 5, 6A/C, 6B, 7F, 8, 9A, 9N, 9V, 10A, 11A, 12F, 13, 14, 15A, 15B, 15C, 16F, 17F, 18B, 18C, 19A, 19F, 20, 21, 22F, 23A, 23B, 23F, 31, 33F, 35F, 37, 38 |
| Magnitude of anionic surface charge | 1, 2, 3, 4, 5, 6A/C, 6B, 7A, 7B, 7F, 8, 9A, 9L, 9N, 9V, 10A, 10F, 11A, 11C, 11F, 12A, 12F, 13, 14, 15A, 15B, 15C, 16F, 17A, 17F, 18A, 18B, 18C, 18F, 19A, 19B, 19C, 19F, 20, 22F, 23F, 27, 29, 31, 32A, 32F, 33B, 33F, 34, 35A, 35B, 37, 43 |
| IPD case-fatality ratio | 1, 2, 3, 4, 5, 6A/C, 6B, 7F, 8, 9A, 9N, 9V, 10A, 11A, 12A, 12F, 13, 14, 15A, 15B, 15C, 16F, 17F, 18C, 19A, 19F, 20, 22F, 23A, 23F, 24F, 31, 33F, 34, 35B, 35F, 38 |

**Figure S1:**
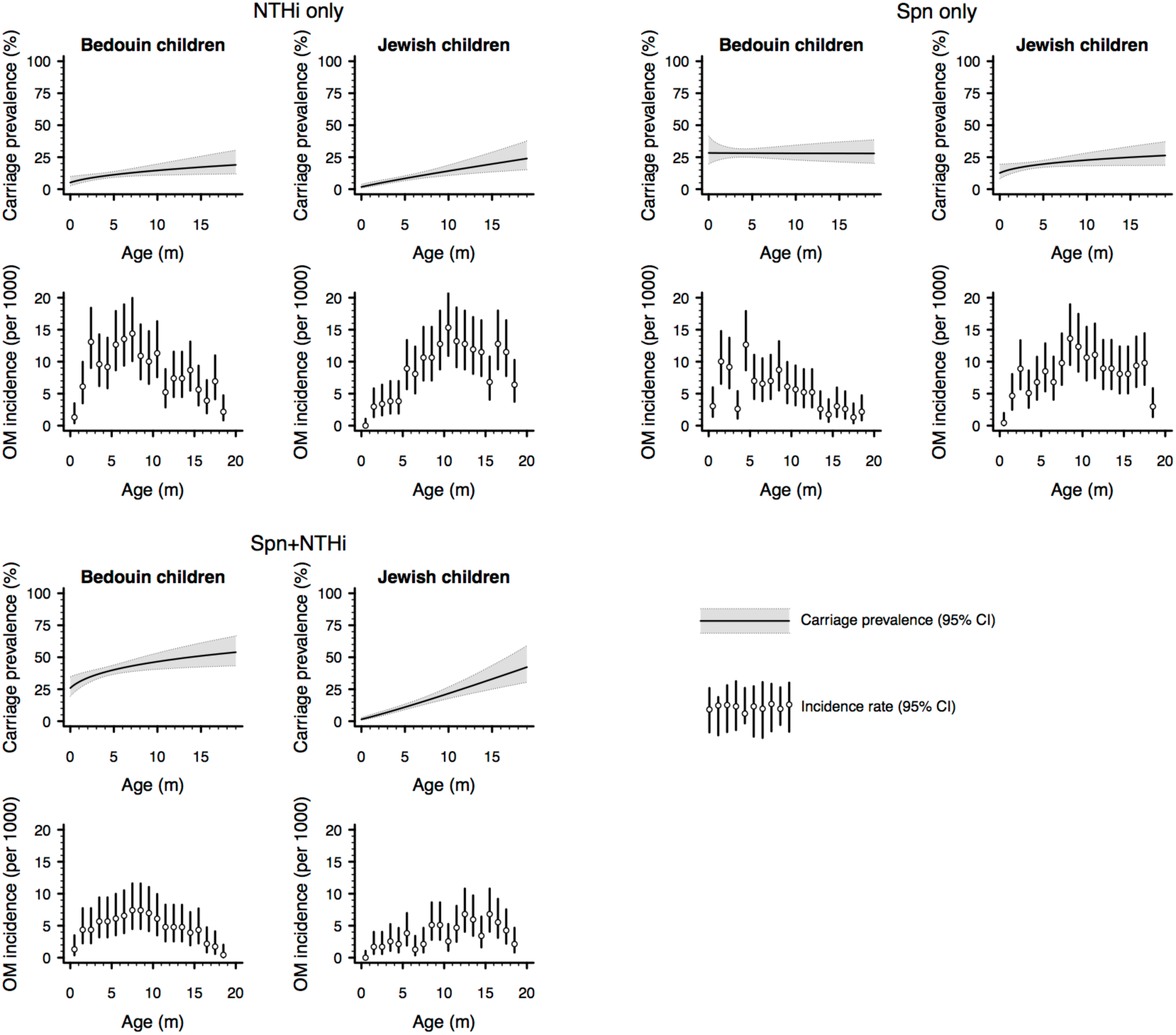
Carriage prevalence and otitis media incidence among Jewish and Bedouin children. We calculated age-specific prevalence of Spn and NTHi carriage by fitting regression models with a log link function to data from scheduled visits at ages 2, 4, 6, 7, 12, and 18 months; we estimated the continuous trend over ages using a log transformation, which we determined was superior to first, second, or third-order polynomial terms via the Bayesian information criterion. We used an exchangeable correlation structure to account for repeated sampling of individual children. Incidence rate analyses were limited to data from 2004 to 2008 for compatibility with the timeframe of the carriage studies, and with previous studies of epidemiological circumstances preceding rollout of PCV7/13 in southern Israel (1, 2). Lines denote 95% credible intervals around estimates.

